# A recursive pathway for isoleucine biosynthesis arises from enzyme promiscuity

**DOI:** 10.1101/2025.08.26.672309

**Authors:** Vittorio Rainaldi, Stefano Donati, Sarah D’Adamo, Nico J. Claassens

**Affiliations:** Laboratory of Microbiology, Wageningen University, Stippeneng 4, 6708 WE, Wageningen, the Netherlands; The Novo Nordisk Foundation Center for Biosustainability, Technical University of Denmark, 2800, Kgs. Lyngby, Denmark; Bioprocess Engineering, Wageningen University, Droevendaalsesteeg 1, 6708 PB, Wageningen, the Netherlands

## Abstract

Enzyme promiscuity can be the starting point for the evolution of new enzymatic activities and pathways. Previously Cotton et al. (2020) identified underground isoleucine biosynthesis routes that can replace the canonical route in *Escherichia coli*, after they deleted the enzymes that catalyze the formation of its precursor 2-ketobutyrate. Using this strain and short-term evolution we identify a new pathway for isoleucine biosynthesis based on the promiscuous activity of the native enzyme acetohydroxyacid synthase II. We demonstrate that this enzyme catalyzes the previously unreported condensation of glyoxylate with pyruvate to generate 2-ketobutyrate in vivo. The gene encoding this enzyme, *ilvG,* is inactivated by a frameshift mutation in the laboratory model strain *E. coli* K-12 MG1655. Its evolutionary reactivation we report here points to a potential natural role in isoleucine biosynthesis in *E. coli.* Isoleucine biosynthesis proceeds with a further condensation step of 2-ketobutyrate with pyruvate, again catalyzed by AHAS, giving the proposed pathway the unusual property of recursivity. The discovered enzyme activity uses glyoxylate and pyruvate as direct central metabolic precursors for isoleucine biosynthesis instead of its canonical ‘indirect’ biosynthesis via the amino acid threonine. Unlike previously discovered underground isoleucine routes by Cotton et al., this route is more likely to play a role in natural isoleucine biosynthesis in *E. coli* due to the use of ubiquitous metabolites and its activity in aerobic conditions. The discovered route further expands the known metabolic space for isoleucine biosynthesis in *E. coli* and potentially other organisms, and could find applications in biotechnological isoleucine production.

## Introduction

Central metabolism includes all the reactions that are necessary to produce building blocks for essential biomass components, such as amino acids for the synthesis of proteins. Amino acid biosynthesis pathways have been the subject of intense scrutiny over many decades to elucidate the biochemical and physiological details of each component of the network (Neidhardt, 1987; Noor et al., 2010; Umbarger, 1978). Despite these efforts, our understanding is incomplete on multiple levels, even for model organisms such as *Escherichia coli.* This is partially explained by the fact that enzymes are often able to accept multiple substrates, a property called promiscuity, leading to overlap and crosstalk between pathways (Noda-Garcia et al., 2018). Promiscuity increases the complexity of metabolism, but it is beneficial in terms of network stability and resilience. Additionally, promiscuous activities can be the starting point for the evolution of new enzymatic activities and pathways (D’Ari & Casadesús, 1998; Khersonsky & Tawfik, 2010).

An emerging method to study enzyme promiscuity and underground metabolism is the use of microbial strains in which auxotrophies for certain key metabolites are created via one or multiple genetic deletions (Kim et al., 2010; Pontrelli et al., 2018; Wenk et al., 2022). Such synthetic auxotrophic strains are often also referred to as biosensors strains or growth-coupled selection strains, as they can ‘sense’ the presence of a certain metabolite or essential building block, with growth as a simple experimental readout. Especially for *E. coli* a wide collection of sensor strains has been built in recent years (Schulz-Mirbach et al., 2025). Metabolites for which these strains are made auxotrophic can be supplied externally in the growth medium to support their growth. When the metabolite is left out of the medium, these strains can be tested with new synthetic routes for the biosynthesis the metabolite, and this method is increasingly used in metabolic engineering of synthetic metabolic routes (Orsi et al., 2021). Alternatively, these sensor strains can be used to identify and potentially evolve latent native routes for biosynthesis of essential metabolites. This approach using synthetic auxotrophic strains has for example led to the identification of novel routes for the essential cofactors coenzyme A and pyridoxal-5′-phosphate biosynthesis in *E. coli* (Kim et al., 2010; Pontrelli et al., 2018).

Recently, an engineered isoleucine auxotroph *E. coli* strain was reported in which all the threonine deaminases were deleted, leading to a lack of the threonine-derived isoleucine precursor 2-ketobutyrate (2KB) (Cotton et al., 2020). Using this strain the authors reported two previously unknown routes for 2KB biosynthesis. The first route is based on the promiscuous O-succinyl-homoserine γ-cleavage of the native cystathionine γ-synthase encoded by *metB*, an essential enzyme in methionine biosynthesis. Another is based on the condensation of propionyl-CoA and formate by 2KB-formate lyase TdcE. Their findings brought the number of known 2KB biosynthesis pathways in nature to nine, of which four have been demonstrated in *E. coli* (Abramsky et al., 1962; Cotton et al., 2020) (Figure 1).

**Figure 1.**
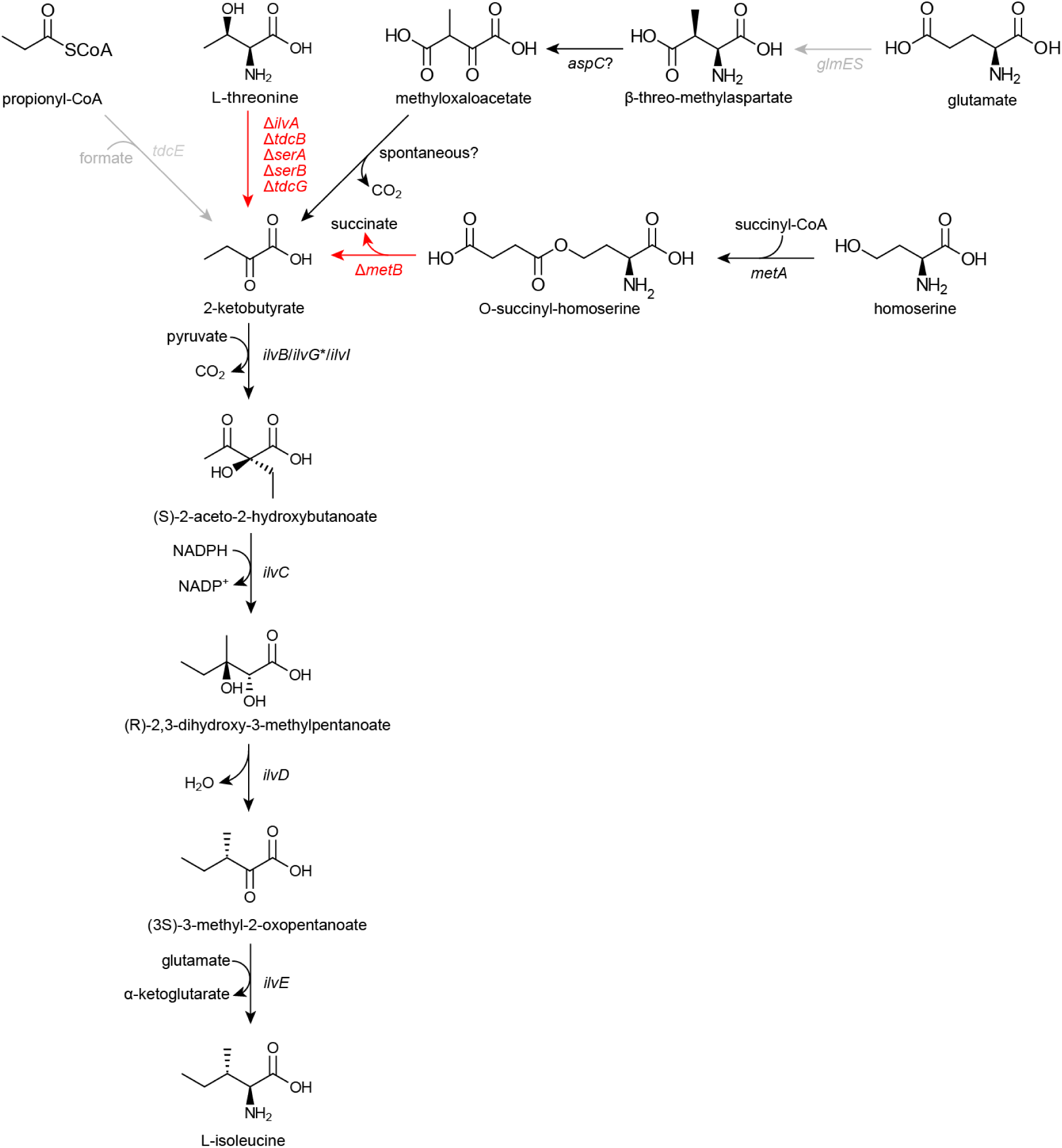
Scheme of the isoleucine/methionine auxotroph strain (IMaux) used in this study as a 2KB biosensor, showing the four previously identified isoleucine biosynthesis pathways in *E. coli*. Red arrows indicate gene deletions, grey arrows indicate reactions that are inactive in the presence of oxygen (in addition, *glmES* is present in some *E. coli* strains but not in *E. coli* MG1655). *ilvG\** indicates that this gene is truncated in *E. coli* MG1655 due to a frameshift and hence inactive

Using a similar isoleucine/2KB auxotrophic *E. coli* strain, we discovered that a key enzyme downstream of 2KB in isoleucine biosynthesis, acetohydroxyacid synthase (AHAS) II (encoded by *ilvG*), also can support the biosynthesis of 2KB. We report that AHAS II is able to accept the smaller 2-keto acid glyoxylate in addition to its canonical substrates pyruvate and 2KB. The condensation of glyoxylate with pyruvate eventually generates 2KB, giving rise to a previously unknown recursive pathway for isoleucine biosynthesis, which we demonstrate with isotopic labeling to elucidate metabolic fluxes. This is the tenth discovered pathway for isoleucine in nature, and provides further insights in the metabolic space and promiscuous networks of amino acid biosynthesis. This shorter isoleucine biosynthesis route may be widespread in nature and can be a potentially attractive pathway for isoleucine production in biotechnology.

## Results

### Characterization of an *E. coli* 2KB biosensor strain

We obtained the Δ5 isoleucine auxotroph strain (SIJ488 *ΔilvA ΔtdcB ΔsdaA ΔsdaB ΔtdcG::kan*) as a kind gift by Dr. Charlie Cotton with the intention of testing alternative 2KB biosynthesis routes (Figure S1). For this reason, we additionally deleted cystathionine γ-synthase (*ΔmetB*), since it was reported that this native enzyme involved in methionine biosynthesis, could also promiscuously support 2KB biosynthesis via O-succinylhomoserine cleavage activity (Figure 1) (Cotton et al., 2020). This deletion in the methionine biosynthesis pathway results in a methionine auxotrophy, so we named the resulting strain isoleucine-methionine auxotroph (IMaux). Another known pathway for 2KB biosynthesis in *E. coli* relies on the reverse activity of 2KB formate lyase (encoded by *tdcE*), however this enzyme requires a glycyl radical and can only operate under anaerobic conditions, and hence its deletion was not required as we only worked under aerobic conditions (Figure 1). A last known pathway, via methylasparate, also did not require a deletion, as this metabolite does not commonly occur in *E. coli* since it requires a glutamate mutase which is not present in *E. coli* MG1655, though it is present in other strains of *E. coli* such as the pathogenic strain O157:H7, where its native role is expected to be glutamate fermentation (Kronen & Berg, 2015), but it may also play a role in isoleucine biosynthesis (Abramsky et al., 1962; Phillips et al., 1972). Additionally, glutamate mutase is an oxygen-sensitive enzyme and it requires the coenzyme B_12_, which *E. coli* cannot synthesize (Fang et al., 2018) and is not present in the M9 minimal medium used in this study.

The IMaux strain cannot grow on glucose or glycerol unless both methionine and isoleucine or 2KB are provided (Figure 2A), confirming the functionality of the sensor strain. Notably, the strain displays a lag phase when supplemented with as little as 0.5 mM of 2KB (Figure 2A), suggesting toxicity of this compound to *E. coli*. The mechanism of this toxicity is unknown, though several hypotheses have been put forward: interference with native metabolism via competitive inhibition (Bisswanger, 1981) due to the structural similarity between 2KB and pyruvate, the role of 2KB as a so-called ‘alarmone’, signaling the shift between aerobic and anaerobic growth (Daniel et al., 1983), and the inhibition of acetolactate synthase (ALS) activity, leading to valine and/or leucine starvation (LaRossa et al., 1987). We reasoned that if the inhibition of ALS activity is responsible for the toxicity effect, supplementation with branched-chain amino acids (BCAA) valine, leucine, and isoleucine should remove the toxicity. Our results indeed demonstrate that 2KB toxicity is mediated by dysregulation in BCAA metabolism, since supplementation removes the lag phase (Figure 2B).

**Figure 2.**
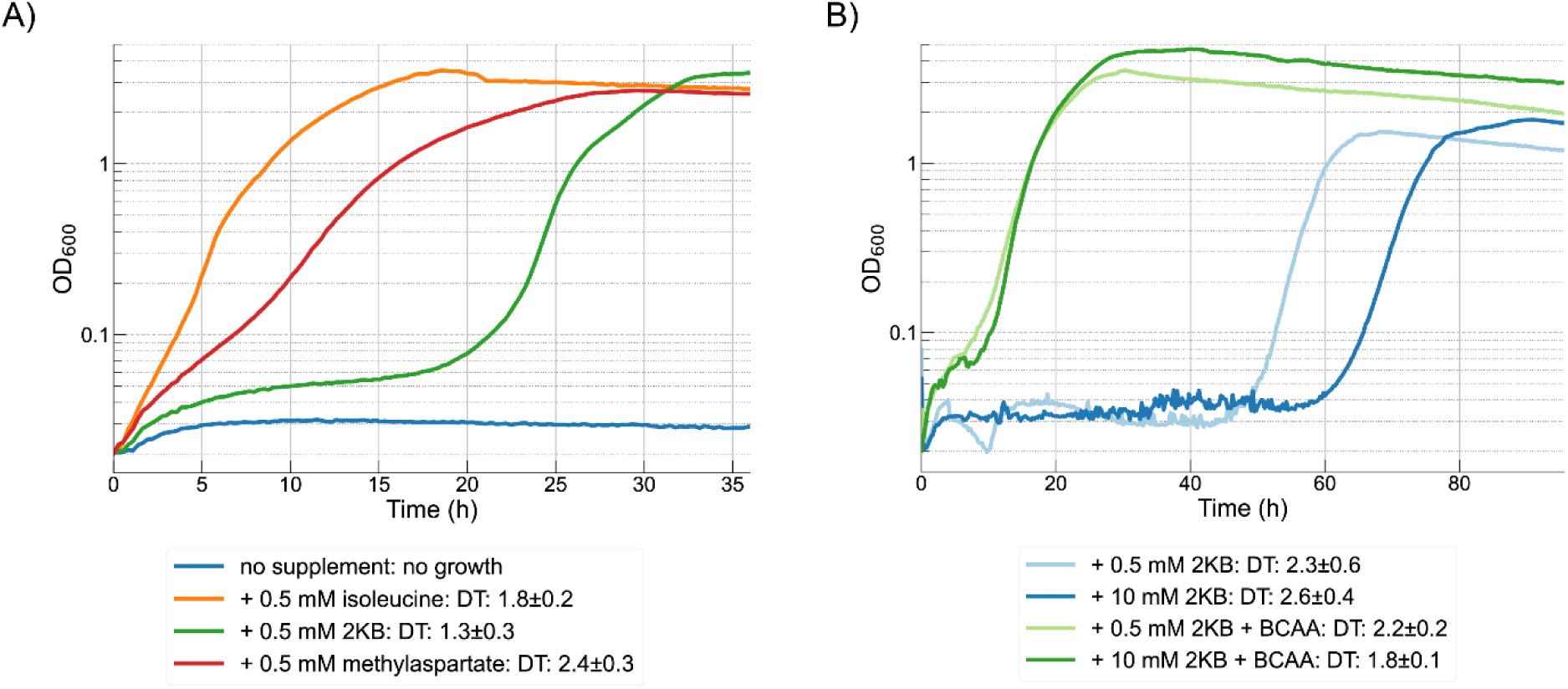
IMaux strain characterization (A) The strain can serve as a sensor for isoleucine, 2KB, or compounds that can be converted to 2KB. Methylaspartate can be converted to 2KB in vivo via promiscuous reactions of native *E. coli* enzymes (B) Supplementation of branched chain amino acids valine, leucine, and isoleucine (BCAA) at 1 mM each relieves 2KB toxicity, indicating that the mechanism involves dysregulation of BCAA biosynthesis. All experiments have been performed in M9 minimal medium with 20 mM glycerol as a main carbon source and 1 mM methionine. Growth curves are the average of at least three technical replicates. DT, doubling time.

### Confirming the methylasparate route for isoleucine biosynthesis in *E. coli*

Next, we tested the strain’s performance as a 2KB biosensor by supplementing methylaspartate, the intermediate that in some *E. coli* strains under anaerobic conditions can be formed from glutamate by glutamate mutase. The formation of 2KB from methylaspartate has been reported in *E. coli* strains W and Crookes (Abramsky et al., 1962; Phillips et al., 1972). This pathway presumably proceeds through transamination to methyloaxaloacetate via the activity of a promiscuous transaminase (most likely aspartate transaminase, AspC), though this activity has not previously been confirmed to the best of our knowledge. The product of this transamination, methyloxaloacetate, is a beta-keto acid which can undergo spontaneous decarboxylation yielding 2KB as a product (Kubala & Martell, 1981), similarly to what is observed for oxaloacetate and pyruvate (Tsai, 1967). Our results show that supplementation of 0.5 mM of methylaspartate fully restores the strain’s growth (Figure 2A), indicating that this sequence of reactions is indeed taking place, and that methylaspartate can be efficiently imported, presumably through one of the native dicarboxylate transporters such as DctA (Unden et al., 2016).

### Discovery of a novel, recursive isoleucine biosynthesis pathway in the 2KB biosensor

We initially intended to test a hypothetical 2KB biosynthesis pathway that requires the condensation of propionyl-CoA and glyoxylate (Figure S1A). For this reason, we performed negative control experiments to test whether the IMaux strain could grow when supplemented with glyoxylate and/or propionate. Unexpectedly, we noticed growth after a lag phase of about 100h when 10 mM glyoxylate was supplemented (data not shown), potentially indicating that this compound can be converted to 2KB via an unknown mechanism.

We isolated and sequenced the resulting strain to confirm the presence and nature of genomic mutations. The only mutation we found was an insertion of two nucleotides in the *ilvG* pseudogene (+AA at position 981) reverting the frameshift that is normally present in the laboratory strains of *E. coli* such as the K-12 derivative MG1655 strain used as a base for these experiments (Gray et al., 1981). This mutation has already been reported, and it is known to lead to the production of the enzyme acetohydroxyacid synthase (AHAS) II (Lawther et al., 1981, 1982). The AHAS II enzyme is one of the three AHAS complexes found in *E. coli.* The AHAS complexes I, II, and III, are encoded respectively by *ilvBN*, *ilvGM*, and *ilvIH*, respectively (Eoyang & Silverman, 1984; Gollop et al., 1989; Hill et al., 1997). Each of these is composed of a larger, catalytic subunit (IlvB, IlvG and IlvI) and a smaller, regulatory subunit (IlvN, IlvM or IlvH). These enzymes were already known for a function in the biosynthesis of isoleucine, but this role is downstream of 2KB biosynthesis, as they condense 2KB with pyruvate into aceto-hydroxbutanoate. This enzyme complex has been studied extensively studied for this role in vitro, and also the promiscuous activity for the condensation of two pyruvate molecules for valine biosynthesis is known (Steinmetz et al., 2010; Vyazmensky et al., 2011).

While IlvG is obviously involved in the isoleucine biosynthesis pathway, it was initially unclear why this mutation would confer the ability to relieve an isoleucine auxotrophy. The known promiscuity of AHAS II led us to hypothesize that glyoxylate might also serve as a substrate instead of the similar keto-acid pyruvate. This leads to a proposed pathway in which AHAS enzymes recursively catalyze two steps isoleucine biosynthesis (Figure 3).

**Figure 3.**
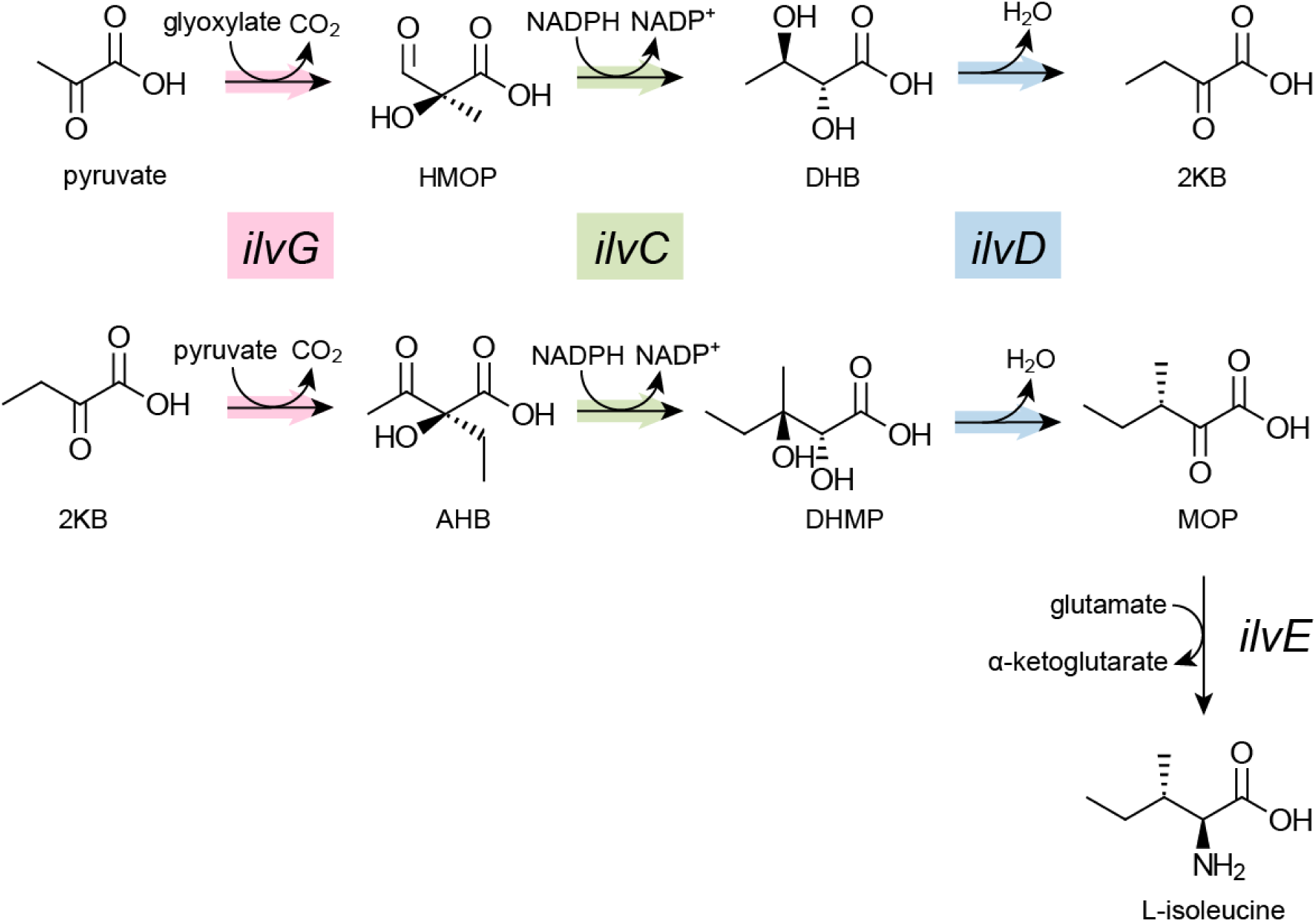
The proposed recursive isoleucine biosynthesis pathway. Pyruvate is condensed with glyoxylate by AHAS II encoded by *ilvG*, then the product of this reaction is reduced and undergoes dehydration to 2KB. The resulting 2KB undergoes the same sequence of reactions to produce the isoleucine precursor (3S)-3-methyl-2-oxopentanoate (MOP), which is further transaminated to isoleucine. AHAS II is the only AHAS enzyme in *E. coli* that can catalyze the first reaction with glyoxylate and pyruvate in vivo, though the condensation of 2-KB with pyruvate can also mediated by AHAS I and AHAS III. Colored arrows indicates that the same enzyme promiscuously catalyzed both reactions. HMOP, 2-hydroxy-2-methyl-3-oxopropanoate; DHB, (R)-2.3-dihydroxybutanoate; 2KB, 2-ketobutyrate; AHB, (S)-2-aceto-2-hydroxybutanoate; DHMP, (R)-2,3-dihydroxy-3-methylpentanoate; MOP, (3S)-3-methyl-2-oxopentanoate.

To verify our hypothesis, we deleted *ilvG* from the strain, which aborts growth on glyoxylate, and confirmed that overexpression of *ilvGM* from an inducible plasmid restores growth. Since *ilvG* is the first gene in the *ilvGMED* operon, a deletion could have disrupted the translation or transcription of the other genes in the transcription unit, making the strain unable to synthesize isoleucine regardless of the supplementation of glyoxylate or 2KB, as *ilvE* and *ilvD* are also essential steps towards isoleucine (Figure 1). To exclude this possibility, we tested both glyoxylate- and 2KB- dependent growth. We confirmed that the uninduced strain was unable to use glyoxylate to synthesize 2KB, while still being able to use 2KB to produce isoleucine (Figure 4A). Based on this result, we concluded that the *ilvG* deletion does not disrupt transcription or translation of the remaining genes in the operon. Moreover, we can conclude that a strain lacking *ilvG* is unable to synthesize 2KB from glyoxylate, meaning that the other AHAS encoding genes *ilvB* and *ilvI* alone cannot substitute for the promiscuous AHAS II activity that is the subject of this paper. This is also implied by the fact that an *ilvG* frameshift reversal mutation was necessary in the evolved strain, and that neither of the counterparts AHAS I and III that are natively present and constitutively expressed were able to support growth on glyoxylate of the 2KB auxotrophic strain.

**Figure 4.**
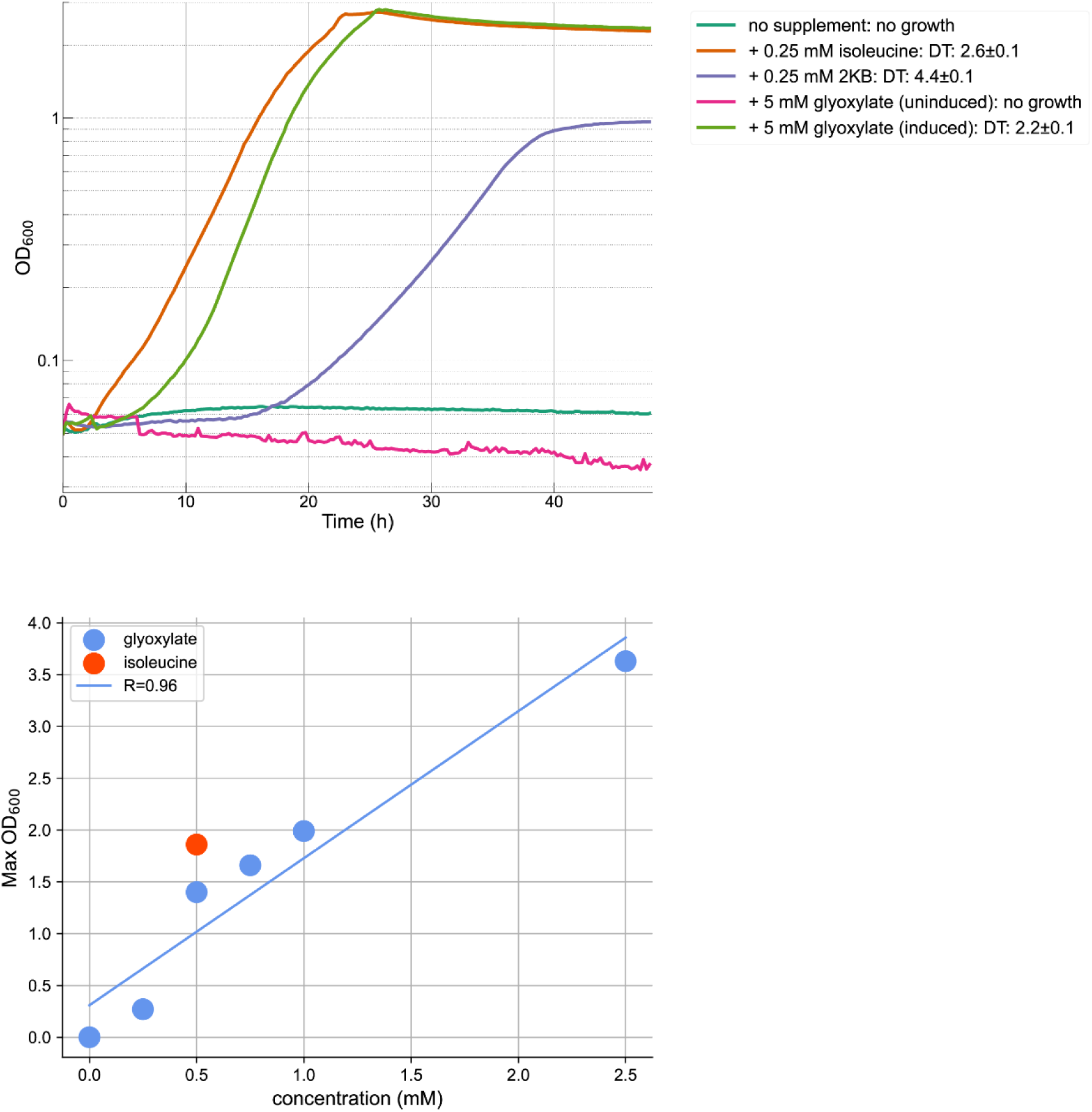
**(A)** Growth of the IMaux *ΔilvG* strain with or without plasmid-based overexpression of *ilvGM*. All strains were grown in M9 with 20 mM glycerol and 1 mM methionine, plus supplemental carbon sources as indicated in the legend. Growth curves are the average of at least three technical replicates. DT, doubling time. **(B)** Max OD reached with different glyoxylate concentrations. Strains were grown in M9 with 20 mM glycerol, 1 mM methionine, and 10 µM cuminic acid as inducer. Blue dots indicate a range of glyoxylate concentrations, orange represents isoleucine (positive control).

As expected, we observed a strict correlation between the quantity of glyoxylate available in the medium and the final biomass density (R=0.96) (Figure 4B).

Next, we used isotopic labeling to confirm our hypothesis that AHAS II produces 2KB from glyoxylate and pyruvate. Based on our proposed route, we expected that if the strains were grown on fully ^13^C-labeled glucose and unlabeled glyoxylate, the fourth carbon (C4) of isoleucine should come from glyoxylate, giving rise to a labeling pattern that is distinct from what arises when supplementing unlabeled 2KB (Figure 5A). The measured labeling of isoleucine confirmed our hypothesis. Notably, in the unlabeled glyoxylate condition, not all of the isoleucine is M+4 labeled, indicating that some of the supplied glyoxylate is feeding into central metabolism (Figure 5B).

**Figure 5.**
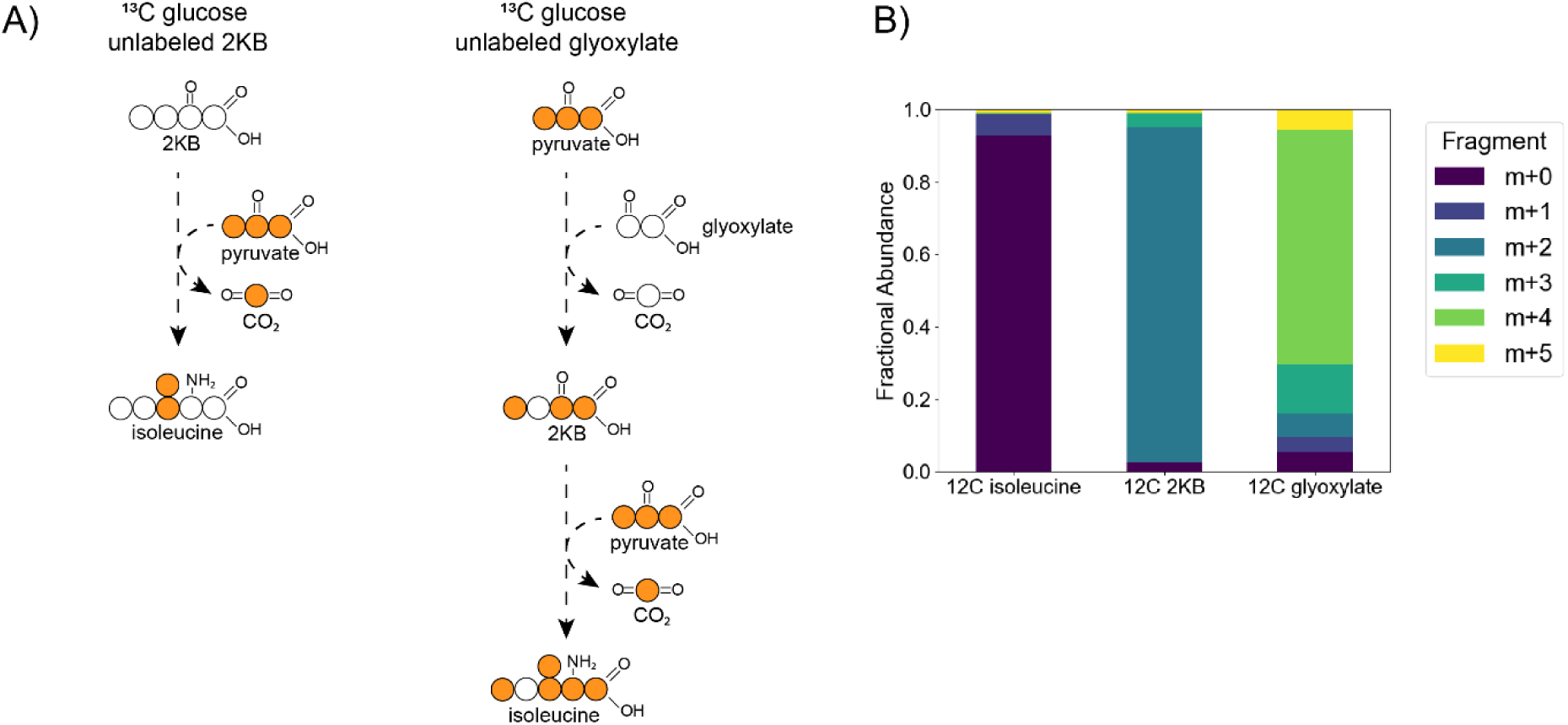
(**A**) Expected isoleucine labeling patterns based on ^13^C glucose and unlabeled 2KB or unlabeled glyoxylate. Orange and white circles represent labeled and unlabeled carbons, respectively. Dashed arrows indicate multiple reactions. (**B**) Observed isoleucine labeling patterns for strains grown on ^13^C glucose supplemented with ^12^C carbon sources as indicated in the figure. The GC-MS derivatization method removes the C1 carbon, leaving only the C2-C6 fragment for analysis. Bars represent the average of three technical replicates.

This could happen via the activity of glyoxylate carboligase (Gcl) and the so-called ‘glycerate route’, or via the activity of malate synthase (AceB or GlcB). Both of these routes should be inactive in glucose minimal medium because of catabolite repression, though it is possible that amino acid starvation leads to their activation mediated by ppGpp (Traxler et al., 2008) or that high glyoxylate concentration supplied in the experiments activate these routes. An intriguing alternative is once again presented by the promiscuous activity of AHAS II: this enzyme could also condense two molecules of glyoxylate to produce tartronate semialdehyde, which could then be reduced to glycerate via IlvC. Glycerate could be assimilated via glycerate kinase (GarK), or undergo dehydration to pyruvate via IlvD.

## Discussion

In this study, we identified a previously unknown, recursive pathway for isoleucine biosynthesis dependent on the promiscuous activity of AHAS II. At least nine pathways for 2KB biosynthesis have been characterized in different organisms (Cotton et al., 2020), however the new route we present here is unique in that it uses an enzyme that is already involved in isoleucine biosynthesis by exploiting a promiscuous activity and producing its own precursor, making the overall pathway recursive, an unusual property in metabolism (Krüsemann et al., 2018; Marcheschi et al., 2012).

AHAS II is thus able to accept a combination of two-, three-, and four-carbon 2- ketoacids as its substrates, leading to multiple roles in BCAA biosynthesis. AHAS II, and also AHAS I and III, also catalyze an essential step in the canonical biosynthesis route of valine, condensing the 2-ketoacid pyruvate with a second molecule of pyruvate to produce acetolactate (Salmon et al., 2006).

Furthermore, unlike several of the other isoleucine biosynthesis routes, the one presented in this study is based on a native *E. coli* enzyme and the central metabolites glyoxylate and pyruvate, pointing to a potential natural role, both in *E. coli* and other bacterial lineages. Indeed, the fact that *ilvG* carries a frameshift mutation in lab isolates of *E. coli* such as MG1655 (Lawther et al., 1981) and BW25113, but not in wild-type isolates from the human gut could point towards the usefulness of this gene in the natural environment. A similar mutation phenotype has been observed in *Salmonella typhimurium* (Burns et al., 1995), providing further evidence for this hypothesis. How and why such *ilvG* frameshift mutations occurred remains unclear: it has been reported that *E. coli* undergoes cycles of isoleucine starvation in minimal medium due to aberrant metabolic regulation when *ilvG* is not present (Andersen et al., 2001). However, since lab strains are generally grown and stored on rich LB agar plates, this effect is presumably reduced or eliminated and may relieve the evolutionary pressure to retain *ilvG*.

The novel identified route could also be beneficial under specific growth regimes, e.g. growth on acetate, as during growth on acetate glyoxylate is produced as a key metabolite in the glyoxylate shunt. Interestingly, it has been shown that *ilvG* expression increases when the *E. coli* K-12 strain MC4100, in which *ilvG* is not natively truncated, is grown on acetate (Oh & Liao, 2000).

Another potential benefit for the route outside *E. coli* could be in photosynthetic or chemolithoautotrophic organisms that generate glyoxylate during CO_2_ fixation using the Calvin cycle. The oxygenase side-activity of the carboxylating enzyme Rubisco, generates phospho-glycolate, which has to be recycled into metabolism via glyoxylate, a key intermediate in all known photorespiration routes (Claassens et al., 2020; Timm et al., 2025). If the described AHAS II activity is present in such organisms, they could use glyoxylate as a building block for isoleucine.

Our results highlight a novel way to synthesize isoleucine based on the common metabolites glyoxylate and pyruvate. We propose that this novel route could be exploited as an isoleucine production pathway in nature and also in industrial isoleucine production (Lu et al., 2024; Zhang et al., 2024). This route, unlike the canonical threonine deaminase route, and O-succinylhomoserine cleavage route, is not linked to other amino acid biosynthesis routes, and requires fewer enzymatic steps from central metabolic intermediates. When isoleucine is made via threonine it is dependent on threonine metabolic regulation including attenuation of the *thr* operon (Lynn et al., 1982; Wenk et al., 2025), allosteric inhibition (Wright & Takahashi, 1977), and product inhibition (Wright & Takahashi, 1977). When isoleucine is made via the O-succinylhomoserine cleavage route it is dependent on methionine and cysteine biosynthesis route and needs lower cysteine levels than normal physiological concentrations. Hence, the 2-AHAS route that starts directly from glyoxylate and pyruvate, independent of other amino acid biosynthesis routes (Cotton et al., 2020) could lead to a more efficient biosynthetic pathway for isoleucine in industrial settings. Furthermore, expression of fewer enzymes results in a lower metabolic burden, which could increase substrate use efficiency and product yield.

We also performed a computational flux balance analysis (FBA) of biomass and product yields for each isoleucine biosynthesis route using both glucose and acetate as feedstocks (Table 1). This analysis shows that the recursive AHAS route in terms of isoleucine yield is as efficient as any of the isoleucine biosynthesis routes known in nature, including the canonical threonine deaminase routes. The predicted biomass yields both from glucose and acetate are even slightly higher if this route is used than the canonical route, confirming it could also likely be a naturally relevant route for the biosynthesis of this cellular building block.

**Table 1.** FBA predictions of theoretical biomass and isoleucine yield for the discovered route and other isoleucine biosynthesis routes in *E. coli*. ^a^ this route requires an anaerobic environment. NA indicates this route is not applicable as anaerobic growth (without electron acceptor) is not feasible.

| **Isoleucine biosynthetic routes** | **Biomass Yield (gCDW/mol glucose)** | **Maximum isoleucine yield (mol/mol glucose )** | **Biomass Yield (gCDW/mol acetate)** | **Maximum isoleucine yield (mol/mol acetate )** |
| --- | --- | --- | --- | --- |
| threonine deaminase (e.g. *ilvA*) | 90.26 | 0.75 | 23.35 | 0.20 |
| recursive AHAS (acetohydroxyacid synthase II, *ilvG*) | 90.46 | 0.75 | 23.37 | 0.20 |
| *O*-succinylhomoserine cleavage (*metB*) | 90.29 | 0.76 | 23.35 | 0.20 |
| 2KB-formate lyase (*tdcE*)<sup>a</sup> | 18.04 | 0.80 | NA | NA |
| glutamate mutase (*glmES*)<sup>a</sup> | 18.11 | 0.44 | NA | NA |

Another potential application of the findings in the paper could be the use as glyoxylate biosensor. Glyoxylate is a key metabolite in several relevant synthetic pathways, e.g. for alternative synthetic CO_2_ fixation routes. A recent work identified and constructed several glyoxylate biosensor strains based on an exhaustive computational analysis of the *E. coli* metabolic network (Orsi et al., 2025). This analysis was unable to identify the recursive AHAS route due to the fact that this promiscuous activity was previously unreported. Our strain could also be used as a glyoxylate (or 2KB) biosensor based on its low biomass demand and robust growth with minimal genetic disruption (see also Figure 4B).

Overall, our findings bring the number of demonstrated isoleucine biosynthesis routes in *E. coli* MG1655 up to five (Figure 6) and the total space of isoleucine biosynthesis routes in nature to 10. This highlights the fact that even well-characterized amino acids biosynthesis pathways in a model organism such as *E. coli* are not fully understood, challenging canonical metabolic knowledge.

**Figure 6.**
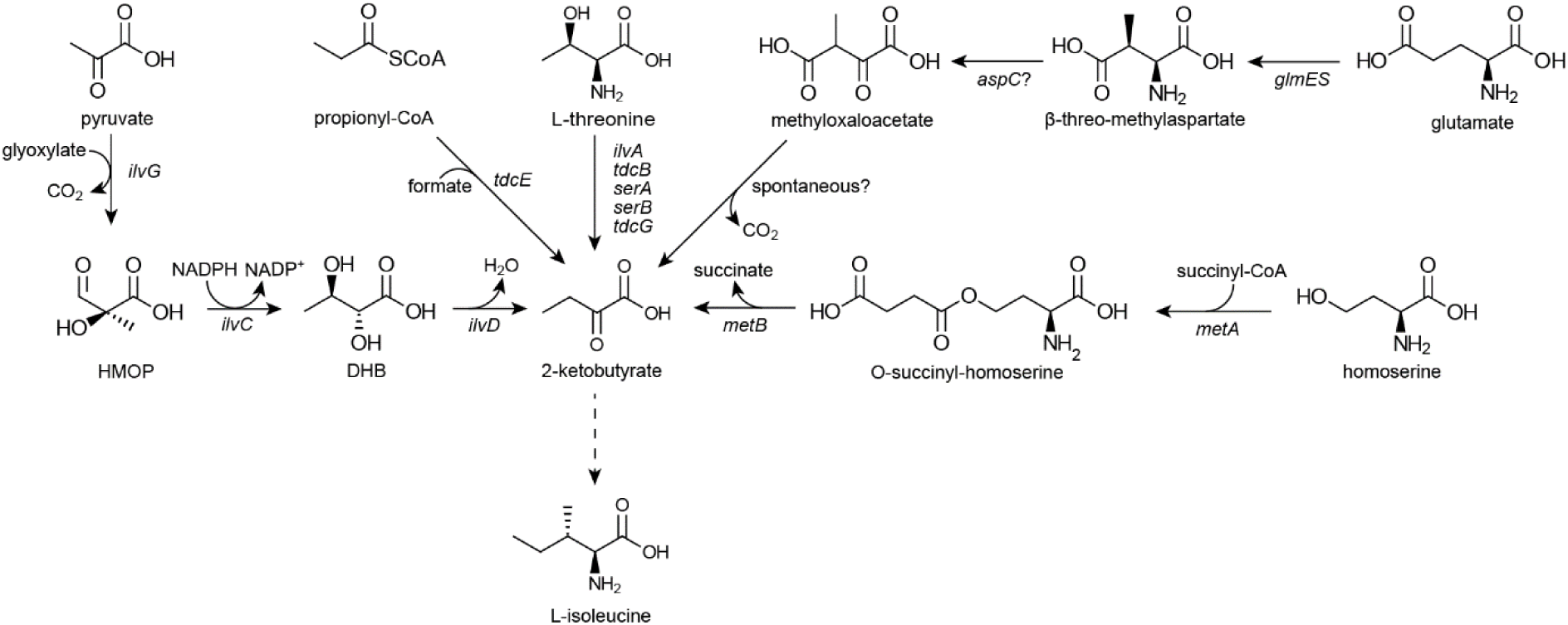
Summary of the demonstrated isoleucine biosynthesis routes in *E. coli*.

## Materials and methods

### Strains and plasmids

All strains used in this study are listed in Table 2. The *E. coli* SIJ488 strain based on K-12 MG1655 (Jensen et al., 2016) was used for the generation of deletion strains.

**Table 2.** Strains and plasmid used in this study.

| Strain name | Genotype | Description | Source |
| --- | --- | --- | --- |
| SIJ488 | *E. coli* K-12 MG1655<br>Tn7:: pAra-exo-beta-<br>gam; pRha-FLP; xylSpm-Iscel | MG1655 derivative<br>with genome<br>integrated<br>recombinase and<br>flippase genes | (Jensen<br>et al.,<br>2016) |
| Δ5 | SIJ488 *ΔilvA ΔtdcB*<br>*ΔsdaA ΔsdaB ΔtdcG::kan* | Threonine deaminase<br>and serine deaminase<br>deletion strain | (Cotton<br>et al.,<br>2020) |
| IMaux | SIJ488 *ΔilvA ΔtdcB*<br>*ΔsdaA ΔsdaB ΔtdcG*<br>*ΔmetB::kan* | Δ5 with O-succinylhomoserine<br>lyase deletion | this<br>study |
| IMaux *ΔilvG* | SIJ488 *ΔilvA ΔtdcB*<br>*ΔsdaA ΔsdaB ΔtdcG*<br>*ΔmetB::kan ΔilvG::cam* | Δ5 with O-succinylhomoserine<br>lyase and AHAS-II<br>deletion | this<br>study |
| Plasmid<br>name | pCym46-ilvGM | cumate-inducible<br>SEVA vector with<br>spectinomycin<br>resistance and p15A<br>ori for the expression<br>of *ilvG* and *ilvM* | this<br>study |

SIJ488 is engineered to carry the gene deletion machinery in its genome (inducible recombinase and flippase). All gene deletions were carried out by successive rounds of λ-Red recombineering using chloramphenicol or kanamycin (pKD3/pKD4) (Datsenko & Wanner, 2000) as described by (Baba et al., 2006). All plasmids are listed in Table 2 and were generated using a Golden Gate protocol as reported previously (Batianis et al., 2020).

### Cultivation conditions

LB medium (1% NaCl, 1% tryptone, 0.5% yeast extract) was used for strain maintenance, generation of deletion strains, and cloning. Antibiotics were used at the following concentrations: chloramphenicol, 25 μg/mL; kanamycin, 50 μg/mL; spectinomycin, 100 μg/mL; apramycin, 50 μg/mL. M9 minimal medium was used for growth experiments (50 mM Na_2_HPO_4_, 20 mM KH_2_PO_4_, 1 mM NaCl, 20 mM NH_4_Cl, 2 mM MgSO_4_,100 μM CaCl2, 134 μM EDTA, 13 μM FeCl_3_ 6H_2_O, 6.2 μM ZnCl_2_, 0.76 μM CuCl_2_ 2H_2_O, 0.42 μM CoCl_2_ 2H_2_O, 1.62 μM H_3_BO_3_, 0.081 μM, MnCl_2_ 4H_2_O) with carbon sources added to the concentrations specified in the text and figures. For growth experiments, overnight cultures were incubated in 3 mL LB medium with appropriate antibiotics. Before inoculation of the experiment, cultures were harvested and washed three times with a saline solution to remove residual carbon sources. Plate reader experiments were inoculated with a starting OD600 of 0.02 using the washed culture. Plate reader experiments were carried out in 96-well microtiter plates (Nunclon Delta Surface, Thermo Fisher Scientific). Each well contained 150 μL of cell culture covered with 50 μL mineral oil (Sigma-Aldrich, Taufkirchen, Germany), to avoid evaporation. A Synergy H1 plate reader (Biotek) was used for incubation (37 °C), shaking, and OD600 measurements. Three cycles of four shaking phases, each of 1 min were used (1. linear shaking at an amplitude of 3 mm, 2. orbital shaking at an amplitude of 3 mm, 3. linear shaking at an amplitude of 2 mm, and 4. orbital shaking at an amplitude of 2 mm). Optical density (OD 600 nm) was measured after each round of shaking (∼12.5 min). Plate reader OD measurements were converted to cuvette values according to the formula ODcuvette = ODplate/0.23. Growth curves were processed in python (https://github.com/he-hai/growth2fig) and represent averages of technical triplicate measurements.

### Whole-genome sequencing

Genomic DNA was extracted using the DNeasy Blood and Tissue kit (QIAGEN) following the manufacturer’s instructions. Genome sequencing was performed by Plasmidsaurus (London, United Kingdom) using the Oxford Nanopore Technology sequencing platform. Analysis of the sequencing data was performed using the breseq pipeline (Barrick et al., 2014) and the SIJ488 reference genome sequence, which is derived from the MG1655 (NC_000913, GenBank). The mutated *ilvG* sequence is provided in the Supplementary file 1.

### Isotopic labeling of proteinogenic amino acids

Strains for isotopic labeling were grown in M9 supplemented with 1 mM methionine using ^13^C labeled glucose and either unlabeled 2KB or unlabeled glyoxylate. The equivalent of 1 mL of OD 1 was harvested, pelleted by centrifugation, and washed twice with sterile distilled water to eliminate leftover traces of medium. The samples were stored at -20 °C and eventually prepared, derivatized and analyzed by GC-MS as previously described (Donati et al., 2023). We considered for further analysis fragments listed in (Long & Antoniewicz, 2019) Table 3, in particular Ile_274. Raw chromatographic data was integrated using SmartPeak (Kutuzova et al., 2020).

Processed data was further corrected for the natural abundance of isotopes in the derivatization agents used for GCMS analysis (Wahl et al., 2004). The data is provided as a Supplementary file 2.

### Flux Balance Analysis

FBA calculations were performed with a custom script on cobrapy version 0.26.3 (Ebrahim et al., 2013), numpy 2.1.1 and pandas 2.2.2. iML1515 was used as *E. coli* genome scale metabolic model for all calculations (Monk et al., 2017). The model was modified by changing the membrane transhydrogenase reaction THD2pp to translocate only one proton, and setting non-growth associated maintenance (ATPM) to 0. Anaerobic reactions PFL, POR5, and OBTFL were also set to zero unless specifically relevant. Oxygen import into the cell (EX_o2_e) was set to zero when modeling anaerobic growth. The promiscuous isoleucine biosynthesis reactions that are the object of this paper, along with the glutamate mutase route, are not part of this model and were added for the relevant calculations.

## Acknowledgements

VR is supported by a VLAG Graduate School Open Round grant. NJC acknowledges funding from the European Union via the ERC Starting Grant FASTFIX. The authors acknowledge Jan Zarzycki and Tobias Erb for fruitful discussions on this work and Tim Althuis for creation of the pCym46 vector.

## Conflict of interest statement

None of the authors declares a conflict of interest.

## Author contributions

VR conceived the work, performed the experimental work, data analysis and wrote the draft manuscript. SD performed the ^13^C -labelling analysis. SA and NJC supervised the work and acquired the funding. All authors helped in revising the manuscript.

---

**Figure S1.**
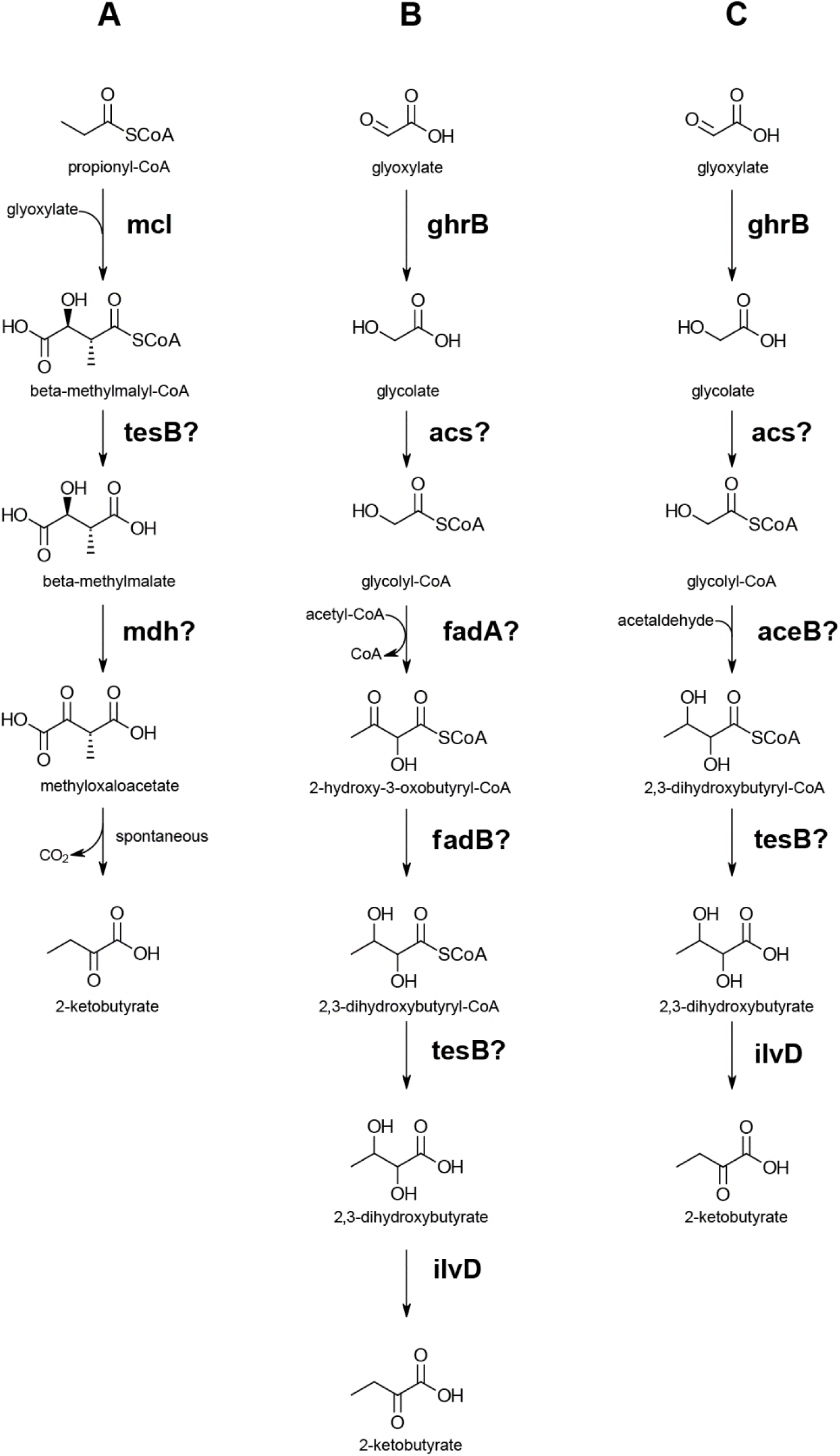
Summary of hypothetical 2KB biosynthesis pathways. Question marks indicate that the indicated reaction has not been demonstrated.

## Notes

### Competing Interest Statement

The authors have declared no competing interest.

